# Investigating the role of non-helpers in group-living thrips

**DOI:** 10.1101/2023.09.08.556834

**Authors:** James D. J. Gilbert

## Abstract

1. Behavioural variation among individuals is a hallmark of cooperative societies, which commonly contain breeders and non-breeders, helpers and non-helpers. In some cases labour is divided, with non-breeders “helping”. Conversely, in some societies subordinate non-breeders often do *not* help. These individuals may be (i) an insurance workforce to ensure continuity of help for breeders when other helpers are lost, (ii) conserving energy while waiting to breed themselves, or (iii) simply of too poor physiological quality either to help or breed.
2. In the Australian Outback, Acacia thrips *Dunatothrips aneurae* (Thysanoptera) glue *Acacia* phyllodes into “domiciles” using silk-like secretions, either alone or cooperatively. Domicile maintenance is important for humidity, so repair can be interpreted as helping. I found that not all females helped to repair experimental damage; some repaired partially or not at all ("non-helpers"). At the same time, some co-foundresses are non- or only partially reproductive, and their role is currently unknown.
3. I first tested the possibility that helping and breeding are divided, with non-helping females breeding, and non-breeders helping. In a lab experiment, I rejected this idea. Experimentally damaged domiciles were typically repaired by reproductive females, and not by non- or partially reproductive individuals.
4. To test whether non-helpers are an insurance workforce, I successively removed repairing females and found that non-helping females did not increase effort as a result. Then, in a field experiment, I tested whether non-helping females were conserving energy while waiting to breed by removing all other females, allowing either a helpful female or a non-helping female to “inherit” her domicile. Isolated like this, non-helpers laid very few eggs compared to helpers or naturally occurring single foundresses, despite having similar ovarian development.
5. My findings show that labour was not divided: reproduction and helping covaried positively, probably depending on individual variation in female quality and intra-domicile competition. Non-helping females were neither an insurance workforce nor conserving energy waiting to breed themselves. They are likely simply of poor quality, freeloading by benefiting from domicile maintenance by others. I hypothesize they are tolerated because of selection for indiscriminate communal brood care in the form of domicile repair.

## INTRODUCTION

Social group members often vary in their contributions to tasks, especially reproduction (Vehrencamp, 1983), but also other tasks like foraging (Johnson, 2010), parental care (Browning et al., 2012), nest defence (Gerber et al., 1988), nest homeostasis (Jandt et al., 2009), and others (see Komdeur, 2006). Costly tasks such as foraging and offspring feeding are often performed by non- or partially reproductive “helpers”, while breeding is performed by one or a few individuals (Koenig & Dickinson, 2004; Solomon & French, 1997). Within some societies, putative “helpers” sometimes do not appear to engage in helping (e.g. chestnut-crowned babblers, Browning et al., 2012; drywood termites, Korb, 2007), especially in small groups (ants, Dornhaus et al., 2008). While this lack of help can often be explained if the helper is waiting to inherit a breeding position (e.g. wasps, Field & Foster, 1999; Leadbeater et al., 2011; lions, Packer et al., 2001), helpers may also engage less in help if they have less genetic stake in current offspring than other individuals (e.g. Galapagos mockingbirds, Curry, 1988); they may function as an "insurance workforce" ensuring continuity of help for breeders in case current helpers are lost (carrion crows, Baglione et al., 2010); or may help in subtle ways such as colony hygiene (Frank et al., 2018; gall-inducing thrips, Turnbull et al., 2012). These explanations implicitly assume that all group members have the same "quality" (i.e. physiological potential for breeding and/or helping). Yet one simple and longstanding explanation for variation in helping behaviour is that group members vary in physiological quality. For individuals of poor quality, helping may constitute the "best of a bad job" (Craig, 1983; Eberhard, 1975). Such individuals would be likely also to provide poor quality help, and thus appear "lazy" compared to individuals of higher physiological potential. There is no unequivocal evidence for this "subfertility hypothesis" (Strassmann & Queller, 1989), although studies of wasps have tended to reject it (Field & Foster, 1999; Leadbeater et al., 2011; Sullivan & Strassmann, 1984).

Acacia thrips, *Dunatothrips aneurae* Mound (Thysanoptera:Phlaeothripidae), live in the arid Australian Outback where they glue Acacia phyllodes (leaflike outgrowths of the stem) together loosely into “domiciles”. Domiciles may be built singly or cooperatively, typically with 2 to ∼7 reproductively mature foundresses (Gilbert & Simpson, 2013; Morris et al., 2002), which are most often sisters but not always (Bono & Crespi, 2008). The factors promoting domicile co-founding in *D. aneurae* are only partially understood, partly because individual roles within groups are unknown. Foundresses do not repel conspecific or heterospecific intruders (Gilbert et al., 2012; Gilbert & Simpson, 2013), although co-founding carries survival advantages especially when invaded by kleptoparasites (Bono & Crespi, 2006). Building and repairing the domicile prevents adults and offspring equally from desiccating, constituting progressive parental care (Gilbert, 2014). No other form of brood care is known. Domicile cohabitation entails reproductive competition, especially in small domiciles, and some females are nonreproductive; their role within the domicile is currently unknown, but they appear to have a negative effect upon other females’ fecundity (Gilbert et al., 2018).

During fieldwork, I noticed that cohabiting female *D. aneurae* vary in their apparent efforts to repair experimental damage to domiciles - some apparently "helping" less than others or not at all (henceforth "non-helpers"). I reasoned that helping by repairing domiciles may be one way for nonreproductive females to gain inclusive fitness - an idea previously suggested by Crespi et al (2004 p.78), perhaps as a means of "paying to stay" (Gaston, 1978). If this is the case, nonreproductives should be the primary responders when domiciles are experimentally damaged, while actively breeding females should be the ones that refrain from helping. If non-helping females are not primarily breeders, they may instead be an insurance workforce for the breeders, repairing only when currently helping females are lost - in which case they should increase their repair effort when any primary-responder females are removed. Alternatively, non-helpers may be conserving energy while they wait to breed themselves, in which case they should increase both breeding and helping effort if experimentally given the opportunity to inherit their domicile. If non-helpers conform to none of these predictions, they may simply be of too poor quality either to breed or help. I therefore asked three questions:

1. Are reproduction and domicile repair divided among foundresses?
2. Are non-helping foundresses an insurance workforce for breeders?
3. Are non-helping foundresses waiting to breed themselves?

To address question 1, I investigated in the lab whether the probability of repairing experimental damage was associated with reproductive status (i.e. ovarian development). It is important to separate helping *probability* from helping *effort*, since the two are not necessarily related (Dornhaus et al., 2008). Thus, in some domiciles, I further quantified repair effort in response to experimental damage. In these domiciles I successively removed females from groups in the order they responded. This allowed me to assess whether repair effort and reproductive status were quantitatively related, and whether repair latency is a good proxy for repair effort. This experiment also allowed me to address question 2 by asking whether initially non-helping females would increase effort after other females were removed (i.e. act as an insurance workforce). Finally, to address question 3, in the field I conducted a removal experiment to test whether initially non-helping females would independently repair the domicile and breed when given the chance - in terms of both potential reproduction (ovarian status) and realized reproduction (eggs laid). I did this by removing all females but one, allowing this focal female, either a helper or a non-helper, to “inherit” her domicile.

## METHODS

### Study organism

*Dunatothrips aneurae* are distributed across the semi-arid interior of Australia, following the distribution of their host plant, *Acacia aneura* (Crespi et al., 2004). They feed on the phyllode surface by piercing and sucking the contents of epidermal cells, as do thrips generally (Chisholm & Lewis, 1984). A "domicile" is founded by one or more mated females (Gilbert & Simpson, 2013) and typically consists of terminal *A. aneura* phyllodes glued together using a silk-like anal secretion (henceforth “silk”). The silk encloses a relatively humid space that prevents death by desiccation in the arid outback, but plays no apparent role in feeding or protection from enemies (Gilbert, 2014). Within this space the females lose their wings and spend the rest of their lives. They outbreed with occasional immigrant males who are attracted to newly built domiciles, and produce a single generation of offspring who disperse after a brief period of sib-mating (Gilbert & Simpson, 2013). Females do not feed outside the domicile and the interior phyllode surface gradually becomes yellow over time, likely indicating feeding activity and a depletion of resources. Population structure in *D. aneurae* is viscous and cofoundresses are most often sisters (Bono & Crespi, 2008), usually having dispersed together by walking, as observed in the lab (the author, unpublished data); coordinated dispersal by flight is presumably unfeasible as the insects are minute and with poor control of flight.

For both lab and field studies, *Dunatothrips aneurae* domiciles were identified and either marked using coloured tape or collected between January and October 2013 at a large patch of *A. aneura* at the Bald Hills paddock (S 30° 57′ 39′′; E 141° 42′ 18′′) near the University of New South Wales Arid Zone Research Station on the Fowlers Gap property, approx. 110km N of Broken Hill, Australia (see Gilbert & Simpson, 2013 for details). Domiciles are not strongly synchronized in developmental stage within or across patches (Gilbert & Simpson, 2013) and the sample contained a mixture of domiciles with and without offspring. I collected domiciles whole, by hand, from the Acacia stems and immediately placed them into ziplock bags inside a cool box, moving them to a laboratory at room temperature within 12 h and maintained using protocols described in Gilbert et al (2013). I examined the domiciles under a dissecting microscope.

### Are reproduction and domicile repair divided among foundresses?

I selected a sample of 48 domiciles containing two or more females. In each domicile, I made a small tear in the wall (roughly 20% of the silk area), folded back the silk, and observed repair behaviour by the foundresses. "Repair" behaviour consisted of stretching single strands of silk between phyllode surfaces. Repair behaviour is obvious, as the female vigorously waves her abdomen before laying down a strand of silk. Typically at least one foundress would begin repairing immediately, within 30 seconds of damage; at this point I gently brushed repairing females on the pronotum with a very small amount of fluorescent paint powder (US Radium Co®), to create a temporary distinctive ID with no noticeable behavioural effects. I continued behavioural observations for 1 h. In these 48 domiciles, I scored females’ behaviour simply as "repair or not". After scoring in this way, all group members were removed and dissected under a microscope (see protocol below) for assessment of ovarian status. I also measured female body size to the nearest 20 µm (pronotum width at 50⨉ magnification).

### Are non-helping foundresses an insurance workforce for breeders?

In a subset of 19 domiciles from those described above, I further quantified repair effort by counting the individual silk strands applied by each female while repairing the experimental damage. For this experiment I selected multi-female domiciles (with 3+ foundresses), in order to test whether females’ repair effort is responsive to removal of other females. If a non-helping female functions mainly as an insurance repairer to protect the domicile’s offspring in case other helpers are lost, her rate of strand deposition should increase once quicker-responding females are removed. After 1 h I removed the first repairer to respond, and observed the behaviour of the remaining female(s) over the subsequent 1 h period. Over successive 1 h periods I removed one-by-one the repairers, ranking them in the sequence that I saw them begin repairing, and counted silk strands laid down by the remaining females. Note that "rank" was determined just by which female began repair first, and I did not distinguish females repairing simultaneously from those repairing sequentially. Only dealate (reproductively mature) individuals were included in experimental observations, as typically only these individuals contribute to repair (Gilbert & Simpson, 2013). Removed individuals were dissected and scored for ovarian development as described below, and their pronotum width measured.

### Are non-helping foundresses waiting to breed themselves?

I used a field removal experiment to address this question (Figure 1). In a sample of field domiciles with 2 or 3 individuals, I first categorised each female as “repairer” or "nonrepairer" using methods described above: I slightly damaged the domicile wall, then observed the behaviour of the inhabitants for 3 h. I used a longer observation period of 3 h under field conditions where observation was more difficult, to be sure to identify nonrepairing females. I did not mark females in the field as the powder marks were difficult to see without a microscope and were quickly lost, but I maintained visual contact and tracked females for the experimental period. Failing to repair within 3 h has meaningful effects because small larvae can die of desiccation in 6 h in the arid outback (Gilbert, 2014). After classifying, I removed all individuals from all domiciles. I then assigned domiciles randomly to treatment groups in which I reinstated to her original domicile either a repairer (R group, n=11), or a nonrepairer (NR group, n=8). In this way I allowed (or forced) this female to “inherit” her domicile. Replacement was completed within 30 minutes of removal. All other females I dissected to assess reproductive status as described below. The remaining female I left in the domicile.

**Figure 1.**
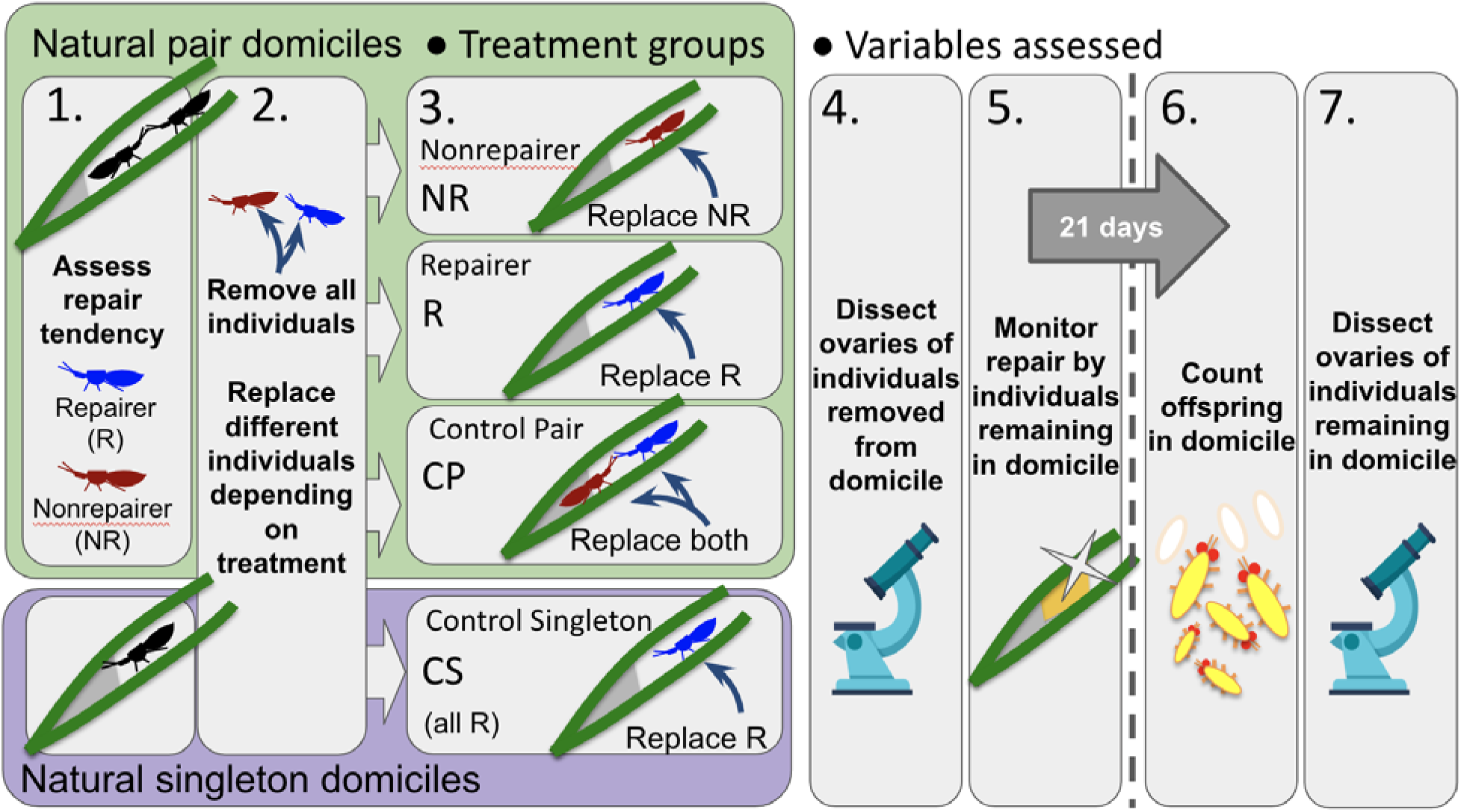
Schematic showing protocol for field removal experiment.

I created two control groups (CS, control singletons, and CP, control pairs) to compare against the two treatment groups (R and NR). The CS group consisted of naturally occurring singly built domiciles (n=46), providing an indication of unmanipulated variation in repair and reproduction by a female that is evidently capable of building a domicile and breeding independently. The CP group consisted of domiciles naturally built by pairs of females (n=9), indicating unmanipulated variation in repair and reproduction where two females were present. In these groups, I performed the same experimental domicile damage before removing and immediately replacing all domicile inhabitants.

After manipulation, I monitored domiciles for 21 days to assess (1) time to repair the domicile, and (2) number of offspring produced. Twenty-one days is the maximum time between domicile completion and egg-laying seen previously in the lab (Gilbert & Simpson, 2013) and I assumed that an individual’s reproduction in the field during that time would be a reliable, standardized snapshot of its egg-laying rate, as a proxy for fecundity. Twenty-one days is also easily long enough for failure to repair a domicile to result in fitness effects for inhabitants (Gilbert, 2014).

### Time to repair domicile

I first established whether my initial classification of females into repairers/nonrepairers was a good proxy for helping effort over a longer period. To do this, I monitored domiciles daily to assess domicile repair. Each day I visually categorised repair as “no repair”, “incomplete repair” or “complete repair”. I also recorded any instances of naturally occurring domicile damage (and its repair), destruction or abandonment.

### Offspring in domicile

After 21 days, all domiciles were collected and taken to the lab to assess the remaining female’s realized reproduction over this period. I counted the number of offspring (eggs and larvae) in each domicile, and dissected the female to examine her ovaries (see below). All individuals remaining in domiciles after the 21 d experimental period were also dissected as described below to assess reproductive status, and the domicile was frozen at -20°C. Owing to time constraints, only a subset of the CS group (n=8) was used for this part of the experiment.

### Assessing ovary status

I assessed ovarian development in all dissected individuals according to protocols originally described in Gilbert et al (2018). Foundresses were killed by immersing in 100% ethanol for 1 minute, and were dissected immediately in water. I recorded the number and volume to the nearest 20µm at 50⨉ magnification (π×length×(width/2)^2^), assuming eggs are an ellipsoid) of any developing oocytes, and also of any mature, chorionated eggs that were ready to lay, and summed the resulting volumes to give a total volume within each ovary. For analysis, oocyte and mature egg volumes were log-transformed after adding the minimum value in the dataset to all values.

### Statistical analysis

All analyses were conducted in R 4.3.0 (R Core Team, 2023). In the lab experiment, I analysed repair effort as a two-step process: first, I analysed factors affecting the probability of a female repairing or not. I used generalized linear mixed models with binomial errors, with "repaired or not" as the binary response and "domicile ID" as a random factor (n=121 females in 39 domiciles). GLMMs were fitted using the lme4 package (Bates, 2010) and tested for overdispersion using the overdisp_fun() R function (GLMM FAQ; Bolker et al; 5 Oct 2022; available at: https://bbolker.github.io/mixedmodels-misc/glmmFAQ.html#overdispersion). In two separate models, I used pronotum width as a predictor, and its interaction with either "volume of developing eggs" or "volume of mature eggs" - fitted separately because these two variables were strongly collinear (Pearson’s correlation, r = 0.95). Models were selected via reverse stepwise model selection and compared using likelihood ratio tests against a Chi-squared distribution, retaining only terms that, when dropped, resulted in a significant reduction in explanatory power (p<0.05).

Then, I analysed factors affecting females’ mean rates of strand deposition, in the subset where this was assessed (n=64 females in 19 domiciles). I used a linear mixed model, with "strands per hour" as the response and "domicile ID" as a random factor. Again, I fitted separate models with pronotum width and either volume of developing oocytes or volume of mature eggs as predictors, plus their two-way interaction. In a separate model, I also sought to confirm whether the quickest responders also repaired at the highest rates, that is, whether the female’s ranked responsiveness within her group was associated with her rate of strand deposition overall. Again I used a linear mixed model, with "strands per hour" as the response and "domicile ID" as a random factor, but with "relative rank" as a predictor. Despite the data consisting of counts, all linear models conformed to assumptions of normality. Models were selected via reverse stepwise model selection as above, and compared using likelihood ratio tests.

I tested whether or not females increase their mean repair effort after the removal of quicker-responding females by splitting the data into a mean repair rate "before" and "after" quicker responding females were removed from the domicile. I use a linear mixed model with "female ID" and "domicile ID" as random factors (n=45 females from 18 domiciles, excluding first responders), and "before/after removal of the quicker-responding female" as a predictor variable.

In the field experiment, to analyse the repair time (in days) and numbers of offspring of the focal female, I used generalised linear models (GLM) with poisson errors and “treatment” as a single predictor variable. Neither model was overdispersed. To avoid multiple comparisons among individual treatment levels, I compared the 4 treatment groups using 3 planned orthogonal contrasts:

1. Control pairs (CP group) against all singletons (pooled mean of CS, R and NR groups).
2. Experimental nonrepairers (NR group) against all other singletons (pooled mean of CS and R groups)
3. Experimental repairers (R group) against control singletons (CS group).

I asked whether R and NR individuals from experimental domiciles differed in reproductive status, both before (i.e. individuals removed immediately) and after the experiment (individuals left in the domicile for 21 days). I used separate linear models of the total volume of developing and mature oocytes (square-root transformed to conform to linear assumptions). As predictor variables, I included repair tendency (R versus NR) and treatment (before versus after), plus their interaction. Again, models were selected using a reverse stepwise procedure and compared using F tests.

The "treatment" term was confounded by the number of females cohabiting in the domicile at the time of removal (“before” individuals all had >1 females in each domicile at the time of removal, while “after” individuals had only 1 individual per domicile). My experiment was unable to separate these two confounded variables, but I was able to shed partial light on the problem by conducting a second analysis that also included individuals in the CS and CP groups. All of these individuals had been “left” for the entire experimental period, but they differed in the number of females in the domicile: 1 for the CS group and >1 for the CP group. For this second analysis, instead of treatment (“removed” versus “left”), I instead used the number of females in the domicile at the time of removal (“1” versus “>1”) as a predictor variable.

## RESULTS

### Are reproduction and repair divided among foundresses?

For lab experiments, I dissected a total of 142 females. Of these, 24 females or 17% (14 repairers and 10 nonrepairers) had no developing oocytes and were assigned an oocyte volume of zero. Among the remaining 118 females dissected, the volume of developing oocytes varied from 6.0×10^-4^ to 3.0×10^-2^ mm^3^. 65 females (46%) had no mature eggs; among the remainder, the volume of chorionated oocytes varied from 1.6×10^-3^ to 2.8×10^-2^ mm^3^. Pronotum width varied from 240 to 360 μm. As in Gilbert et al (2018), pronotum width was not correlated with volume of developing oocytes or volume of mature oocytes. The volume of domiciles used in lab experiments ranged from 24.1-934 mm^3^ (median 258 mm^3^).

Of 142 females from 48 domiciles, 109 (77%) were observed repairing (29 began within 1 hour of damage, 80 began only after removal of one or more females) while 33 (23%) were not observed repairing. In 7 domiciles more than 1 female began repairing damage immediately. A simple X^2^ test revealed that females were more likely to participate in repair if they had developing or mature eggs than if they did not (X^2^ = 4.32, df = 2, p = 0.02, Figure 2a). In the GLMM analyses accounting for nest ID, a female’s probability of repairing experimental damage in the lab depended only upon her volume of developing oocytes (GLMM, X^2^=3.92, df=1, p=0.04), or, in a separate model, mature eggs (GLMM, X^2^=4.57, df=1, p=0.03, Figure 2b); no other predictors were significant (all p > 0.05).

**Figure 2.**
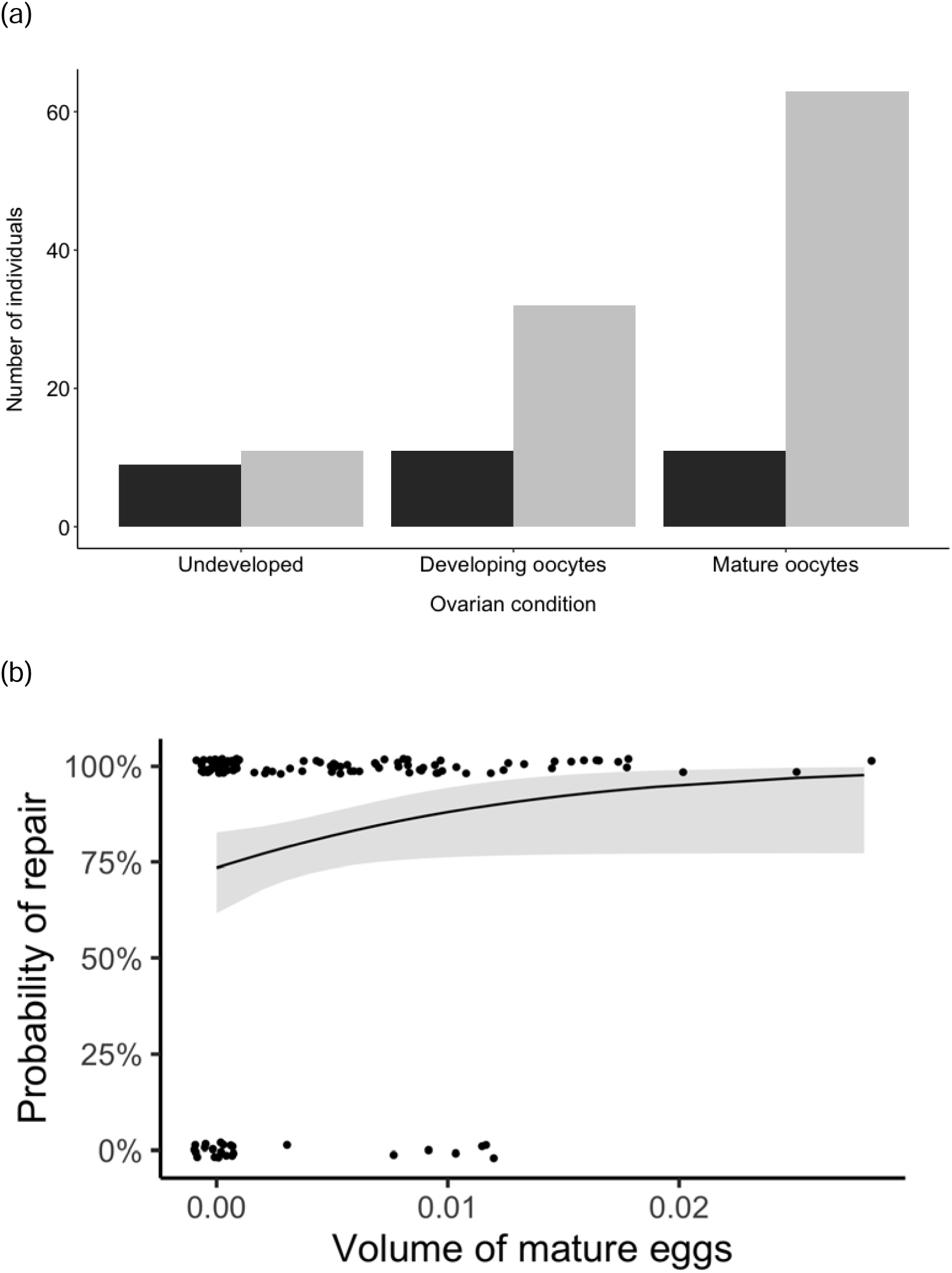
(a) Frequency of participation in domicile repair (black = no repair, grey = repair) by female *D. aneurae* in different reproductive condition; (b) Probability of repair in relation to volume of mature oocytes in females. Fitted line is from the binomial GLMM (see text for details).

### Are non-helping females an insurance workforce for breeders?

In the sequential removal experiment, removing a responder had no statistically significant effect upon the repair effort made by other females (linear mixed model, female ID and domicile ID as random effects, dropping "before/after removal of the quicker-responding female", X^2^=0.22, DF=1, p=0.22, Figure 3). For subsequent analyses of repair rate, I therefore pooled repair rates for each female before and after removing quicker-responding female(s) and used only "domicile ID" as a random effect.

**Figure 3.**
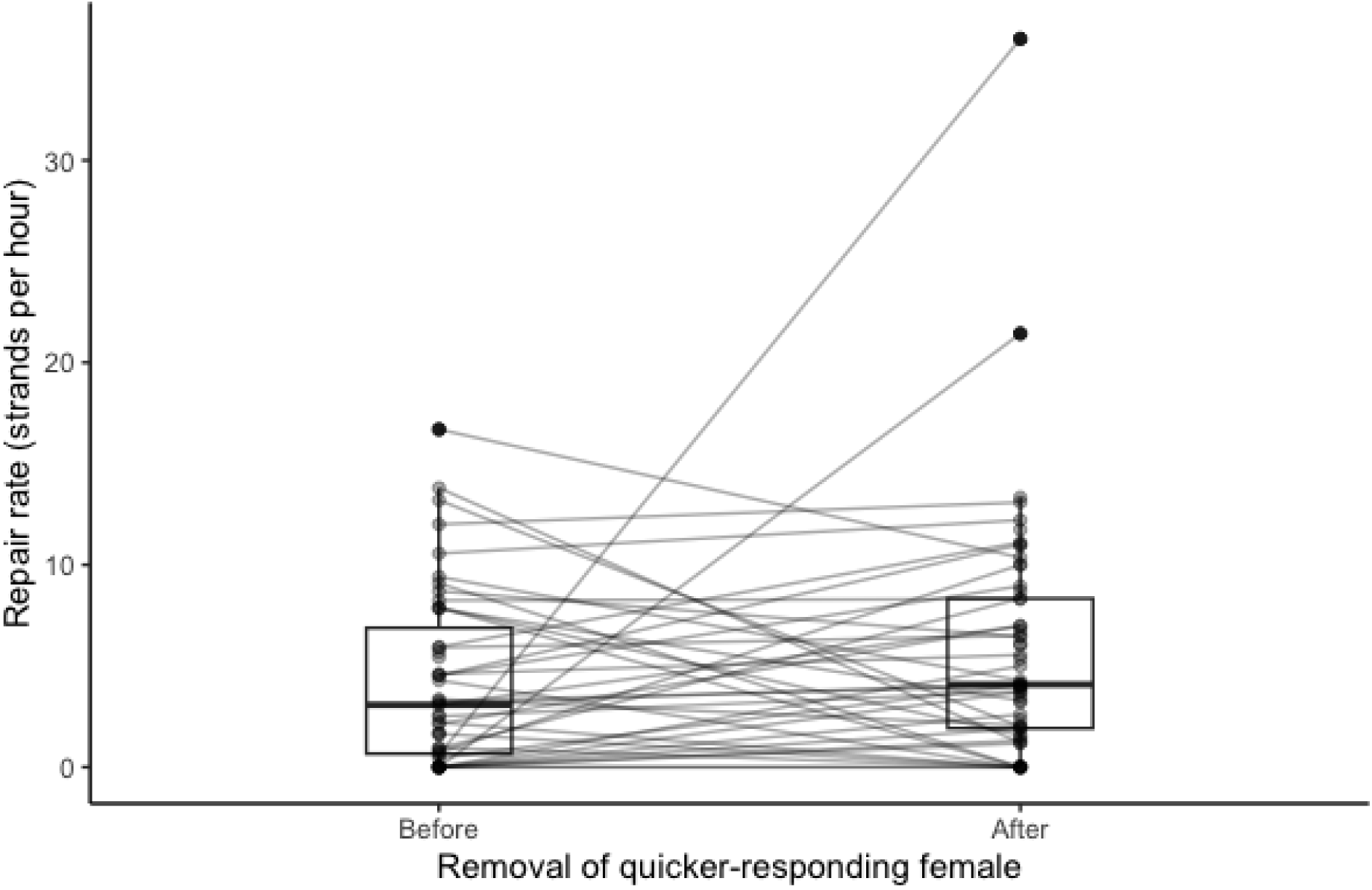
Repair rate (strands deposited per hour) of a focal female, before vs. after removal of the quicker-responding female. Lines join measurements for each individual female.

Thus pooled, total repair rate was positively associated with females’ oocyte volume (linear mixed model with domicile ID as a random effect, =:^2^=4.51, df=1, p=0.03) although not her volume of mature eggs (=:^2^=2.36, df=1, p=0.12). Females with below average volumes of developing oocytes repaired at a mean rate of c. 3-5 strands per hour, whereas females with above-average volumes of developing oocytes repaired at 6-10 strands per hour (Figure 4a). Pronotum width was not associated with repair rate in either model.

**Figure 4.**
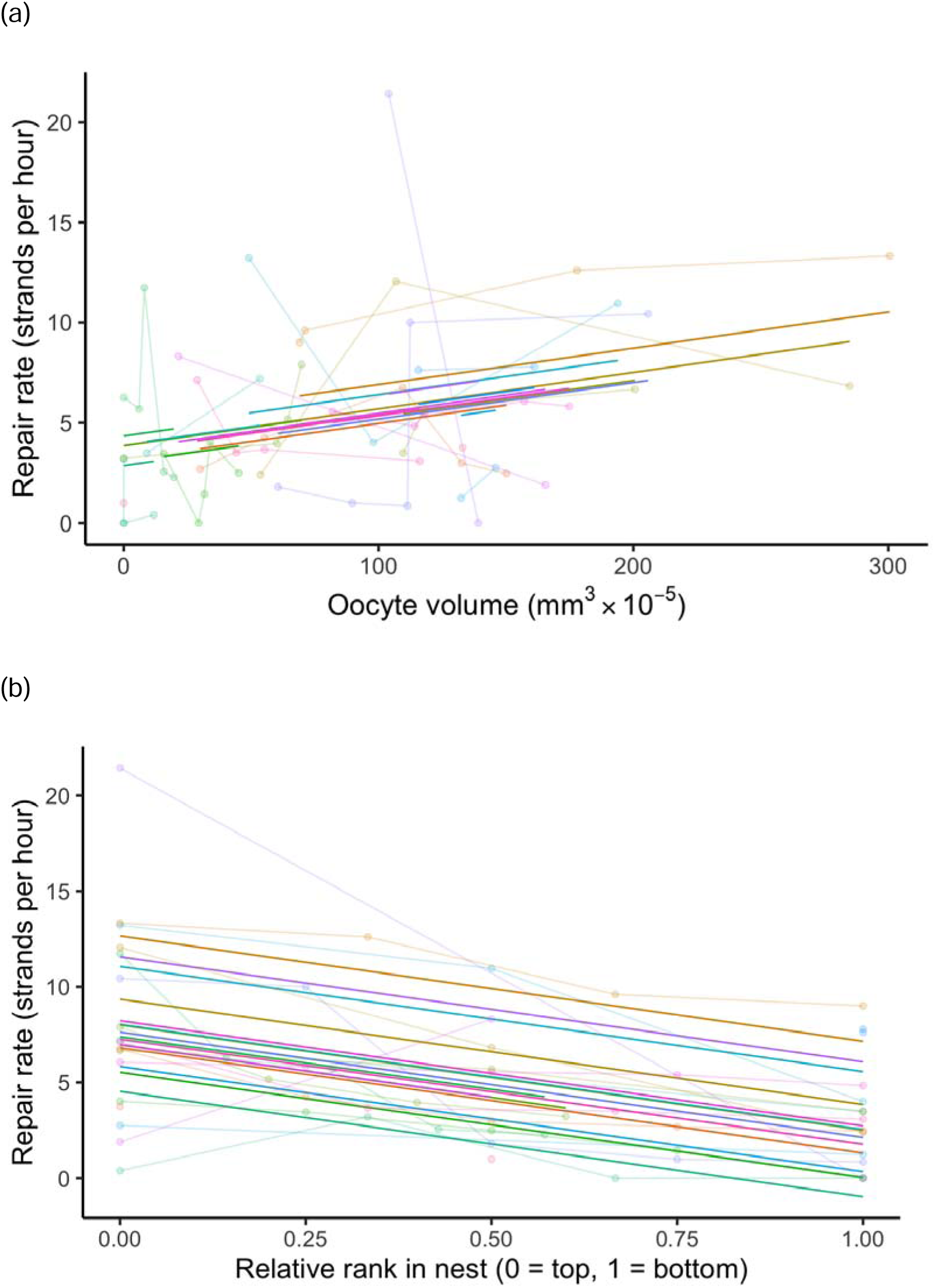
(a) Repair rate (strands laid per hour) in relation to oocyte volume; (b) repair rate (strands deposited per hour) in relation to relative responsiveness rank within domicile. Thin coloured lines connect females within a domicile; heavy coloured lines are estimates for each domicile from respective LMMs (see text for details).

In a separate model, a female’s mean repair rate over the duration of the experiment was predicted by her ranked latency to respond within her group (linear mixed model of overall repair rate with domicile ID as random effect, dropping "female rank", =:^2^=23.10, df=1, p<0.0001; Figure 4a). That is to say, females that responded quickly also repaired at the highest average rates, and vice versa. Oocyte volume did not predict repair rank, i.e. latency to begin helping, within domiciles (GLMM with relative rank as response and nest ID as random effect, binomial errors, removing "oocyte volume" from model, □:^2^=1.04, df=1, p=0.31), and neither did the volume of mature eggs (=:^2^=1.80, df=1, p=0.18)

### Are non-helping foundresses waiting to breed themselves?

In the field experiment, of 21 domiciles with 2 females, I categorised 8 as (R, R), 11 as (NR, R) and 2 as (NR, NR). Of 9 domiciles with 3 females, 1 was (R, R, R), 3 were (R, R, NR) and 5 were (R, NR, NR). From these, I assigned 5 ⨉ (R, NR), 2 ⨉ (R, R), 1 ⨉ (R, R, NR) and 1 ⨉ (R, NR, NR) to the "Control Pair" group. The remainder were assigned to R or NR groups. Out of the 46 singleton domiciles in the CS group, all 8 whose repair activity I assessed in the field were repairers. Among repairers, latency to repair varied from <30 seconds to 126 m within the 3 h observation period.

### Time to repair domicile

There were significant differences in repair time among groups (GLM, poisson errors, Δdeviance=43.76, Δdf=3, p=0.004), driven by the NR group (Figure 5a). The NR group took 4 d to complete repair (range 2 – 7 d). In contrast, Control Pairs, Control Singletons and R group females repaired their domiciles in 2 d, with some completed by the next day (range 1-4 d [CP], 1-2 d [CS], 1-2 d [R]). Contrast 3 was significant (NR group against pooled mean of CS and R groups; z=-3.143, p=0.002), indicating that nonrepairers differed from other singleton females. No other contrasts were significant. There were also no differences in the frequency of domiciles destroyed by wind or otherwise abandoned during the experimental period (*X*^2^=0.625, p=0.884).

**Figure 5.**
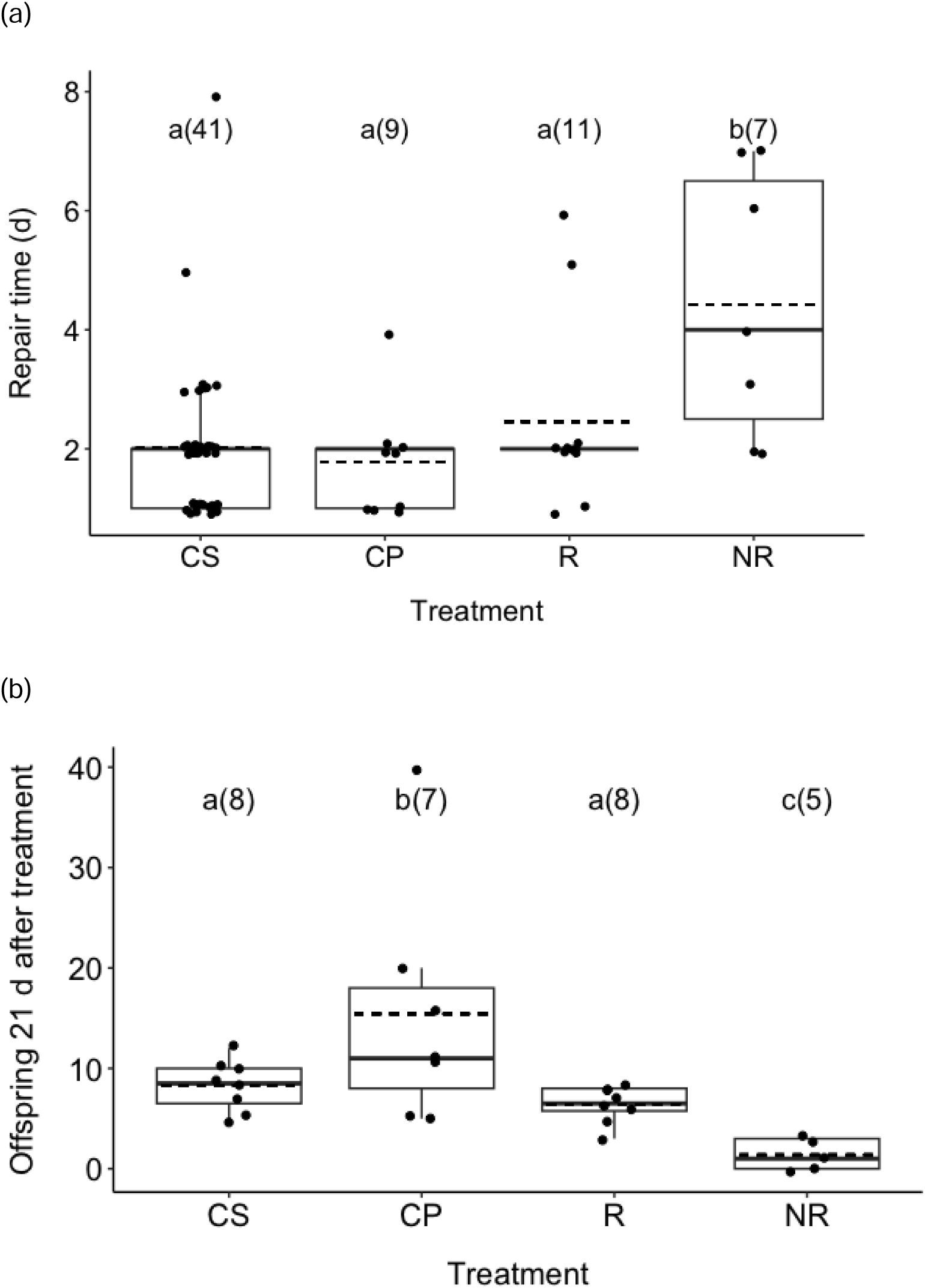
(a) Median time to repair damage in *D. aneurae* domiciles from different treatment groups. (b) Median number of eggs in the domicile 21 d after treatment. A small amount of jitter has been applied to separate overlapping points. Bars with different letters are statistically different. Sample sizes (domiciles) are given in parentheses. Dashed line shows mean.

### Offspring in domicile

Despite data being counts of offspring, a GLM with poisson errors was overdispersed and I found the data conformed to standard linear assumptions, so I fit a linear model. The number of offspring after 21 d was different among groups (F_2, 23_ = 8.67, p < 0.0001; Figure 5b). Notably, the NR group had substantially fewer offspring than all other groups (Contrast 3, NR group against pooled mean of CS and R groups; z = 3.487, p<0.001). Control Pairs had more offspring than all other groups (Contrast 1, CP against pooled mean of CS, R and NR; z = -3.89, p<0.001). Experimental repairers were similar to control singletons in productivity (Contrast 2, R against CS; z = -1.13, p=0.27).

### Reproductive status

In the field experiment, five dissected females (3 repairers, 2 nonrepairers) had no developing oocytes. Among the remainder (n=82), the volume of developing oocytes varied from 5.3×10^-4^ to 3.4×10^-2^ mm^3^. Thirty-six had no mature eggs, and among the remainder (n=51) mature egg volume varied from 2.0×10^-3^ to 3.2×10^-2^ mm^3^.

There were no significant differences in volume of developing oocytes or of mature eggs, whether among experimental treatments or repair tendencies (linear models, all p>0.05).

## DISCUSSION

In multi-female domiciles of *D. aneurae*, helpers were more likely than non-helpers to be reproducers (Figure 2). Reproducers were also *better* helpers: females with more developing oocytes put in more effort while repairing (Figure 4a). Quick-to-respond repairers were also good repairers, and *vice versa*, both in the lab (Figure 5a) and in the field (Figure 5a), making repair latency a good proxy for overall helping effort. In the field, good repairers laid more eggs (Figure 5b) although they did not have more oocytes or mature eggs when dissected.

Non-helpers were not an insurance workforce for breeders, since females did not adjust their help in response to changes in group composition (Figure 3). Neither were they waiting to breed themselves, since they did not increase reproduction when other females were removed (Figure 5b). Below, I argue that nonreproductive non-helpers are likely of too poor quality either to substantially help or breed, and that it is possible they remain in the domicile as parasites whose few offspring benefit from care by others (i.e. domicile maintenance). Finally, I hypothesize they may be tolerated by breeders because of selection for communal, indiscriminate care.

### Reproduction and domicile repair were not divided among foundresses

Contrary to my prediction, I found that reproduction and helping effort were positively correlated in this species. Non-breeding non-helpers may comprise 25% of social groups in some cooperative societies (Teunissen et al., 2020) so the concept of "lazy" helpers is not new (Korb, 2007). Repair and reproduction may be positively associated if, for example, only high quality individuals (those that can acquire significant resources) are capable either of repairing or reproducing. This could come about if females vary significantly in quality, and thus resource acquisition rather than resource allocation determines their capacity both to breed and repair (van Noordwijk & de Jong, 1986). However, it is important not to assume *a priori* that silk production is costly; its cost has yet to be demonstrated and testing for this cost should now be a priority for research into this species.

Alternatively, a link between repair and reproduction may be mechanistic; for example, the silk used to bind domiciles together may be the same as the glue used to stick eggs to phyllode surfaces within domiciles, such that only reproductive females are capable of performing either task. A similar link might be found, for example, in honeybees where only individuals producing queen mandibular pheromone (ie. queens) are capable either of attracting males or of eliciting retinue behaviour in workers (Jarriault & Mercer, 2012). The physiological basis of glue/silk production should now be a focus for researchers interested in group living thrips.

### Non-helping foundresses were not an insurance workforce for breeders

What is the role of those group members who breed and help partially or not at all? Non-helpers seemed to carry no detectable benefits to domiciles in terms of integrity: in the field experiment, domiciles built by control pairs versus singletons were repaired equally quickly (Figure 5a) and suffered similar rates of wind damage or abandonment. I rejected the idea that these females function as insurance helpers, since they did not increase effort when quicker-responding females were removed. Instead, females helped at a relatively fixed rate which was correlated with their reproductive status (Figure 4a) and also with how quickly they began helping (Figure 4b). Consistent differences in subordinate contributions to helping have been shown across social contexts in, for example, social spiders, at least temporarily (Settepani et al., 2013) and meerkats (English et al., 2010). On the other hand, helpers in carrion crows (Baglione et al., 2010) and molerats (Mooney et al., 2015) appear to show a degree of responsiveness to social context.

The presence of non-helping females did not appear to translate into any particular reproductive benefit for breeders (as do helpers in some cooperative breeding societies, e.g. Nelson-Flower et al., 2013), since females had no more developing oocytes in cofounded domiciles than in singleton domiciles. In previous studies, females sharing domiciles with nonreproductives actually had *fewer* developing oocytes (Gilbert et al., 2018). This suggests that non-helping females may not gain indirect, kin-selected benefits from whatever little help they can provide.

It is still possible these females may help breeders in subtle ways. For example, subordinates may act via “load lightening”, extending longevity of breeders, even though direct effects upon current breeding may not be obvious (Russell et al., 2007). Bono and Crespi (2006) found that cooperating females enjoyed a survival advantage over singletons when invaded by kleptoparasitic thrips. Additionally, "non-helpers" may actively participate at the building stage of the domicile, or may contribute to maintenance of middens (Gilbert & Simpson, 2013). In other social insects, nonreproductive individuals can have subtle but important effects upon colony function; for example, “soldiers” help to combat pathogens in thrips (Turnbull et al., 2012) and ants (Frank et al., 2018).

### Non-helping foundresses were not waiting to breed themselves

I also rejected the idea that non-helping nonreproductive females were maximising their direct fitness by conserving energy for breeding themselves later, since they did not take the opportunity to increase their reproduction when it was experimentally offered to them. To my knowledge, this is the first study in which nonbreeding group members failed to increase their reproduction when given this opportunity (see e.g. Field & Foster, 1999; Leadbeater et al., 2011; Rehan et al., 2014; Smith et al., 2009). Note that this finding also rejects the idea that non-reproductives are reproductively suppressed by the presence of breeders, as in e.g. meerkats (Young et al., 2006), molerats (Bennett et al., 2022) and cichlid fish (Heg, 2008). It is conceivable that these females may have “come online” more than 21 days after manipulation, and subsequently either dispersed and bred independently or bred within their existing domicile. However, dispersal is unlikely because all mature female *D. aneurae* lose their wings (Gilbert & Simpson, 2013) and there is no evidence yet for multiple generations or domicile reuse (see Gilbert & Simpson, 2013) as in other social species (e.g. Rehan et al., 2014), probably due to depletion of plant cell food resources within domiciles after one generation.

One possibility is that these females lacked sufficient physiological resources either to breed independently or to substantially help. Variation in both breeding and helping may arise from significant variation in quality (genetic or developmental) (Grinsted & Bilde, 2013), and/or from variation in condition (Parthasarathy et al., 2022) that may come about via within-group competition (e.g. Young et al., 2006). Reproductive failure is not uncommon in some social species at rates of up to 50% (Brahma et al., 2018; Grinsted & Bilde, 2013; Salomon et al., 2008). Reproductive competition in *D. aneurae* occurs both in smaller domiciles and in domiciles containing more females (Gilbert et al., 2018), as well as those which are occupied by inquilines (Gilbert et al., 2012). It is likely that poor quality females also lose out in competition for feeding; little is known about females’ positioning, although they do move frequently within domiciles (pers. obs.). Females with low competitive ability may fail to secure the nutrition required for oogenesis (Wheeler, 1996) or may need to resorb oocytes (Boggs & Ross, 1993) after performing presumably costly tasks such as domicile repair, even at low intensity (note that "non-helpers" did eventually repair their domiciles, although taking twice as long as repairer females; Figure 5a). This may explain why non-helpers ended up laying much fewer eggs than repairers (Figure 5b), despite having similar volumes of developing/mature eggs within ovaries.

For a poor quality, non-helping female, there may still be direct benefits of remaining in (or joining) an established domicile, because the small number of offspring she does produce will be more likely to survive in a communally maintained domicile than if she had attempted to breed alone. Because domicile repair benefits all females and offspring indiscriminately, her offspring will benefit from maintenance performed by other domicile residents (Jones et al. 2007). Such females may effectively be parasitic, owing to their neutral (this study) or negative effect on breeders’ reproduction (Gilbert et al., 2018).

Why poor quality, apparently costly *D. aneurae* females are tolerated by reproductive foundresses remains an open question, as originally raised in Gilbert et al (2018), until both their role and relatedness within the domicile can be clarified. One attractive possibility is suggested by models of social evolution based on "assured fitness returns" (Gadagkar, 1990; Queller, 1989) in which any amount of help (or offspring) that a female contributes to a social group stands a greater chance of being translated into successful descendants (direct or indirect) than if she attempted to breed alone. In particular, the "fostering model" of (Jones et al., 2007) was originally developed for social spiders and has been used to explain the evolution of inter-species brood care in mixed-species spider associations (Grinsted et al., 2012). The model applies to species with a prolonged period of offspring dependency on parental care, where individual adults are likely to die before offspring are independent from care. Under these circumstances adults will be selected to breed communally so that offspring will receive "foster care" whether or not a given individual dies. *D. aneurae* satisfies many of these conditions: larvae develop slowly, die quickly when exposed, and are reliant on adults for domicile maintenance for the entire developmental period (Gilbert, 2014). The model requires care to be independent of relatedness, a condition ancestral in social spiders (Samuk & Avilés, 2013) and that also applies to domicile repair in *D. aneurae*, which benefits all inhabitants regardless of relatedness.

Selection for communal, indiscriminate care will naturally create an opportunity for a degree of freeloading by parasitic individuals, whether conspecifics (potentially explaining the tolerance of poor quality female *D. aneurae*, as in this study) or heterospecifics such as the inquiline *Akainothrips francisi*, which are tolerated by residents despite imposing reproductive costs upon them, and who benefit from residents’ maintenance of the domicile (Gilbert et al., 2012).

In this study I have rejected some potential explanations for the behaviour of non-helping female *D. aneurae*. An investigation of more subtle roles and the costs of silk production, coupled with a detailed assessment of relatedness within domiciles, are now the main priorities in this species.

## ACKNOWLEDGEMENTS

I thank Prof. S. J. Simpson for mentorship, guidance and use of his lab and facilities, Dr K Leggett and G. and V. Dowling for helpful discussions; help during fieldwork and use of facilities, Dr LA Mound and Dr A Wells for assistance with fieldwork and helpful discussions; Dr T Flower, Dr M Nelson-Flower, Dr LE Browning, Dr L Grinsted and Dr A Russell for helpful discussions; and Dr P. Bierzychudek for supplying micronized paint. This study was funded by the European Community’s Seventh Framework Programme (FP7/2007–2013) under grant agreement no: PIOF-GA-2011–299506.

## CONFLICT OF INTEREST

I declare no conflict of interest.

## STATEMENT ON INCLUSION

My study does not include scientists based in the country where the study was carried out. I recognise that more could have been done to engage local scientists with my research as the project developed, particularly those from the First Nations of Australia, and to embed my research within the national context and research priorities. I am endeavouring to address these caveats in future research.

## DATA AVAILABILITY

All data will be archived in Data Dryad upon publication.

## Notes

### Competing Interest Statement

The authors have declared no competing interest.

